# Causal network perturbation analysis identifies known and novel type-2 diabetes driver genes

**DOI:** 10.1101/2024.05.22.595431

**Authors:** Yue Zhao, Hash Brown Taha, Ansarullah, Kailiang Qian, Parveen Kumar, J. Matthew Mahoney, Hao He, Joshy George, Sheng Li

## Abstract

The molecular pathogenesis of type 2 diabetes mellitus (T2DM) involves environmental and genetic factors that remain poorly understood. Increased insulin resistance leads to pancreatic β-islet cell exhaustion and failure, leading to more severe T2DM. As such, targeting β-cell dysfunction is an attractive pathway for T2DM treatment. Single-cell RNA sequencing can help identify differentially expressed genes (DEGs) that may contribute to disease-associated pathophysiology, but cannot capture the full spectrum of disease pathophysiology, as some genes that are not differentially expressed may still play important roles in disease progression or mechanisms. As such, we investigated single-cell gene expression changes in β-cells from healthy (C57BL/6J) and diabetic (NZO/HlLtJ) mice fed with normal or high-fat, high-sugar diet (HFHS) across the spectrum of T2DM development (prediabetic, mild or severely diabetic) using an innovative integration of the causal network perturbation assessment (ssNPA) framework with meta-cell transcriptome analysis to identify driver genes for T2DM. By generating a reference causal network and conducting *in silico* perturbations, we identified three classes of genes implicated in T2DM pathophysiology: (1) DEGs that were not perturbed (e.g, *Igf2bp2* strongly linked to β cell dysfunction and glucose regulation), (2) perturbed non-DEGs (e.g., *Glp1r* a therapeutic target of modern diabetes drugs) that were perturbed, and (3) DEGs that were perturbed (e.g., *Pdx1* a master β cell transcription factor), and validated some of the targets using the KOMP database. Together, these results show that each analytic layer captures complementary mechanisms underlying T2DM severity, and their synthesis provides a more complete view of the disease pathophysiology.

## Introduction

Type 2 diabetes mellitus (T2DM) comprises more than 90% of all diabetes cases, impacting approximately 6.8% (537 million individuals) of the global population in the year 2021^1,2^. T2DM is caused by a complex interplay between environmental and genetic factors. However, while some of the major environmental factors (i.e., diet and physical activity) are known, the genetic bases of T2DM remain poorly understood. The endocrine portion of the pancreas is constituted by highly specialized hormone-secreting entities known as islets of Langerhans. The islets comprise α, β, delta, PP, and ghrelin cells that secrete glucagon, insulin, somatostatin, pancreatic polypeptide, and ghrelin, respectively^3^. The onset of full-blown T2DM occurs when β cells suffer from failure and exhaustion leading to the loss of their capacity to secrete appropriate amounts of insulin in response to elevated blood glucose levels as a result of insulin resistance in peripheral tissues.

Current research insights on T2DM are limited due to the variability in environmental and genetic factors among diverse populations. Additionally, there is a notable gap in long-term clinical trials that comprehensively assess the efficacy of emerging treatments across different stages of the disease. In recent years, single-cell RNA sequencing (scRNA-seq) has proven critical for investigating comprehensive gene expression profiles, revealing the presence of heterogeneous gene expression patterns, even within the same cell-types^4^. Furthermore, diverse phenotypes of pancreatic β cells have been observed within a single islet^5–7^. The application of scRNA-seq has significantly contributed to our understanding of β-cell maturation, β-cell heterogeneity, β-cell failure, and β-cell function in both healthy and diseased states. However, scRNA cannot capture the full spectrum of disease pathophysiology, as some genes that are not differentially expressed may still play important roles in disease progression or mechanisms. Additionally, no study to date has delineated the gene changes associated with clinical transitions across T2DM’s spectrum by comparing prediabetic, mildly, and severely diabetic stages.

To answer this gap, we used two mouse lines (C57BL/6J and NZO/HILTJ) that were kept on a standard or high-fat high-sugar (HFHS) diet regimen to create the following phenotypes: healthy/non-diabetic (C56BL/6J on standard diet), prediabetic (C56BL/6J on HFHS diet), mildly diabetic (NZO/HILTJ on standard diet) or severely diabetic (NZO/HlLtJ on HFHS diet). At the end of the treatment, we performed scRNA on isolated β-cells to make a comparison between healthy and prediabetic states; prediabetic and severely diabetic states and mildly diabetic and severely diabetic states (**Figure 1A-B**). Additionally, since conventional differential expression analyses do not effectively detect certain type of disruptions in gene networks, we performed a causal network perturbation assessment (ssNPA) framework^8^ in combination with a meta-cell transcriptome analysis (meta-ssNPA) (**Figure 1C)**. By generating a reference causal network and conducting *in silico* perturbations, we identified three classes of genes implicated in T2DM pathophysiology: (1) non-perturbed DEGs, (2) perturbed non-DEGs and (3) perturbed DEGs, and validated our candidates using the Knockout Mouse Phenotyping (KOMP) Project database.

**Figure 1.**
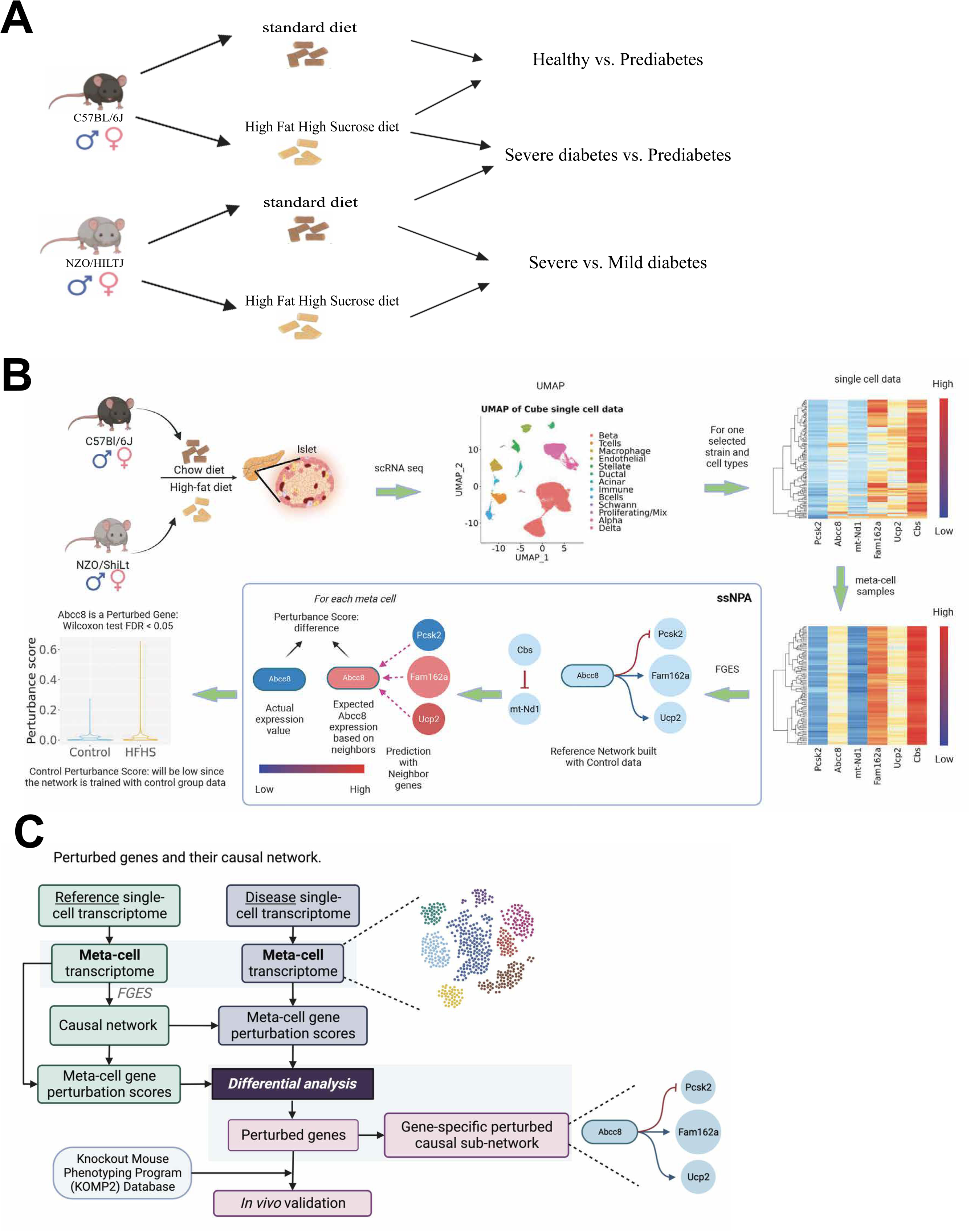
Overview of the meta-ssNPA framework and experimental design. **A)** Experimental setup showing comparisons across the T2DM continuum. C57BL/6J mice on a standard diet were compared to those on a high-fat, high-sucrose (HFHS) diet to model healthy versus prediabetic states, while NZO/HlLtJ mice on standard and HFHS diets represented severe versus prediabetic and severe versus mild diabetic conditions, respectively. **B)** Schematic representation of the single-cell network perturbation analysis (ssNPA) pipeline. Single-cell RNA sequencing data from islets of control and HFHS-fed mice were processed to construct cell-type-specific networks, compute perturbation scores, and identify gene-specific causal relationships using the FGES algorithm. **C)** The meta-ssNPA framework integrates reference and disease single-cell transcriptomes into a unified meta-cell network, quantifies gene perturbation scores, and performs differential analyses to identify perturbed genes and their causal subnetworks. Selected genes were further validated using the Knockout Mouse Phenotyping Program (KOMP) database.

Our findings reveal that integrating differential expression, perturbation, and shared DEG-perturbed analyses provides a comprehensive understanding of T2DM severity mechanisms. The DEG-only layer highlighted genes such as *Igf2bp2*^9^ that regulate insulin signaling and glucose metabolism, while the perturbed-only layer identified *Glp1r* across all three comparisons, a clinically relevant therapeutic target modulating β cell function^10^. The perturbed-DEG layer captured core regulatory genes like *Pdx1*^11^, essential for insulin synthesis and secretion. Together, these complementary layers illustrate how distinct molecular processes converge to drive β cell dysfunction and metabolic imbalance in T2DM.

## Material & Methods

### Animal studies and islet isolation

In strict adherence to the standards set by the Association for Assessment and Accreditation of Laboratory Animal Care, our facility at The Jackson Laboratory has upheld the care and treatment of mice. We acquired male and female mice from two distinct strains: C57BL/6J (B6; RRID:IMSR_JAX:000664), and NZO/HlLtJ (NZO; RRID:IMSR_JAX:002105), starting at the age of four weeks. The mice were provided with two types of diets from Research Diets: a high-fat, high-sucrose diet (HFHS, comprising 44% kcal from fat and 1360 kcal from sucrose; formula D19070208) and a control diet (10% kcal from fat and devoid of sucrose; formula D19072203), both containing equal fiber content. The diets were given *ad libitum* starting from the age of six weeks. At the age of fifteen weeks, islet isolation was performed, and mice were euthanized through cervical dislocation and the common bile duct at the Sphincter of Oddi was clamped. Collagenase solution (three mL of a solution containing collagenase P (5 units/ml) and DNaseI (1mg/ml) in Hank’s Balanced Salt Solution (HBSS)) was inserted into the bile duct proximal to the final bifurcation leading to the liver to inflate the pancreas. We then removed the pancreas for digestion by the collagenase solution at 37°C for 40 minutes. Post-digestion, samples were agitated for ten seconds, diluted it with 10 mL of HBSS, and centrifuged for 3 minutes at 300 RPM. After two washes with HBSS, the pellet was resuspended in 5 mL HBSS, and handpicked islets were collected using a clean petri dish containing HBSS. The islets were then transferred to 24-well plates with 1 mL of the warmed media (containing RPMI 1640, 10% FBS, glutamine, and HEPES) and incubated overnight at 37°C. After overnight incubation islets were centrifuged and the supernatant was discarded. Finally, islets were resuspended in a 1-2 mL of preheated (37°C) StemPro Accutase dissociation solution (Fisher Scientific, cat # A1110501). The cell suspension was gently pipetted for 30 seconds to facilitate the dissociation of islets until the media appeared translucent and there were no visible clumps, usually within 2-5 minutes. Following dissociation, 2-3 mL of RPMI complete medium was added, and the cell suspension was filtered through a 20 μm strainer. The cells were then centrifuged at 230 RCF for 3 minutes, the supernatant was removed, and the cells were resuspended in 2-3 mL of RPMI complete medium.

### Preprocessing of Single Cell RNA-Seq

We processed raw fastq reads from Illumina sequencing for scRNA-Seq by aligning them to the mouse reference genome (mm10/GRCm38) using 10X Cell Ranger version 6.1.1 with standard settings. To demultiplex the strain of origin from samples containing mixed strains, we employed demuxlet version 2 (https://github.com/statgen/popscle)^12^, utilizing known genomic variations from VCF files obtained from the Sanger Mouse Genomes Project^13^. We focused on sites that were biallelic and varied among our two strains (B6, and NZO). We ran demuxlet on scRNA-seq data with specific parameters “--α 0.0 -- α 0.5 --tag-group CB --tag-UMI UB --field GT”, accounting for all cells identified by Cell Ranger.

### Quality Control and Filtering of Single Cell RNA-Seq Data

Post-processing, we applied quality control measures to the feature count matrices generated by Cell Ranger. Single cells with fewer than 500 (islet) genes, more than 20% (islet) mitochondrial transcripts, or over 50% ribosomal transcripts were excluded. Additionally, genes not detected in at least three single cells per sequenced library were also omitted. To correct for potential ambient RNA contamination in islet samples, we utilized decontX from the celda V1.10.0 package, adhering to developer guidelines on GitHub. The SCDS V1.10.0 package was deployed to eliminate cell doublets, using default settings.

### Clustering and Identification of Cell Types in Single Cell RNA-Seq

Following the filtration and quality control, single cells/nuclei underwent normalization and were clustered using Seurat version 4.1, with batch variations across libraries corrected by harmony version 0.1.0. Gene expression data from single cells were normalized based on library size and log-transformed. Dimensionality was reduced using principal component analysis (PCA) on the 2,500 most variable genes, and these principal components (PCs) underwent batch correction using harmony. The batch-corrected PCs were then used for Louvain-based clustering, with the resolution parameter adjusted between 0.1 and 1 according to the dataset specifics.

We identified differential marker genes for various groups via the Wilcoxon rank-sum test or MAST within the Seurat package, employing an false discovery rate (FDR) cutoff of 0.1 and a fold change cutoff of 1.5. Cluster-specific genes aided in assigning cell types. Independently, cell type assignment was also performed unbiasedly using the SingleR package, comparing each single-cell transcriptome to reference transcriptome profiles of known cell types. For formal differential gene expression tests, we created pseudobulks within cell types by aggregating gene expression counts across all cells of a given type from a single mouse, applying DESeq2 on the pseudobulks. This approach using negative binomial modeling with pseudobulks derived from single-cell transcriptomic data is noted for its efficacy and accuracy in differential expression testing in single-cell contexts^14^. Low-expression genes have been filtered out the OGFSC algorithm^15^.

### Refine the existing Single Sample Network Perturbation Assessment (ssNPA) method

There are many limitations that still exist for the current ssNPA framework^8^. First, the framework is not well suited to handle single cell data. The single cell data have sparsity issues where a lot of genes have zero expressions in many cells. This violates the assumption of the linear model ssNPA is using and thus hinders accurate prediction performance. We tackle this problem by a simple meta-cell idea^16^. We randomly pick several cells from the same cell type and average the gene expressions as a new meta-cell sample. This approach maintains the relative level of all gene expression in the cell (high expression genes are still higher and lower expression genes are still lower in the meta-cells) while removing the zeros. Secondly, the current ssNPA framework uses a simple criterion to filter out Perturbed genes: as long as the average Perturbance Score is higher in the test group than the control group, the gene is annotated as a Perturbed Gene. This approach does not consider the random noise in the Perturbance Score, thus induces false positive results. We refined this process by applying Wilcoxon test^17^ onto test and control group. Perturbance Score and use a FDR < 0.05 as the threshold to filter the Perturbed Genes where statistical significance is introduced in the comparison. We performed one sided Wilcoxon test with the alternative hypothesis that the control group Perturbance Score < test group Perturbance Score.

### Workflow and experimental design to detect perturbed genes via meta-cell local reference causal network

Based on the discussion above, we propose the following meta-ssNPA workflow (**Figure 1**): the scRNA-seq data from various sample groups underwent standard processing using a single-cell pipeline to identify distinct cell types. Subsequently, meta-cell transcriptome data was generated for each of these identified cell types. Using the control group samples as a reference dataset, perturbation scores were computed for all genes within a given gene network, employing the ssNPA framework. Perturbance scores were calculated by comparing the network predictions derived from the reference network and the test group data, with the expectation of distinguishing between the test and control groups. Then, we visualize all samples perturbance score with t-distributed stochastic neighbor embedding (t-SNE) and compare them against real group ID (test or control). After verifying that the test and control group are separated well in visualization indicating the Perturbance Scores can be used to distinguish the two groups. The Wilcoxon tests^17^ were applied to filter out genes exhibiting significant perturbations, and finally, pathway analysis was conducted for further interpretation of the findings. The genes with a significant adjusted p-value (< 0.05) in the Wilcoxon test between test and control perturbance score is defined as Perturbed Genes.

We combined a meta-cell transcriptome analysis with the ssNPA framework to identify the genes that are perturbed in the test condition (HFHS diet) compared to that of the control using a causal network^8^. The causal network is computed using a Fast Greedy Equivalent Search (FGES) algorithm^18^ on a directed acyclic graph (DAG), where the nodes are genes, and the directed edges are the causal relationships between the genes. The FGES algorithm iteratively adds an edge if the addition decreases the Bayesian Information Criteria (BIC), where the BIC is calculated based on a linear model where the expression of a gene is predicted by fitting a linear model with the expression levels of its parents, spouses, and children (Markov Blanket) in the current network as the predictors. The single-cell gene expression data from the mouse given the standard diet is used to construct the causal network. The difference between the predicted and actual values (i.e., perturbation score) is used to identify genes essential for T2DM pathogenesis. Since, unlike the standard differential expression analysis, this method considers the interactions between the genes and their neighbors, several genes identified in our analysis could not have been identified otherwise. Importantly, the expression level of perturbed genes can be similar between test and control groups, but the interactions between them and their neighbor genes vary.

### DEG analysis

To identify DEGs associated with diet, we applied the Wilcoxon test on scRNA-seq expression data between control and test group for β cells using the same code in^17^. Genes with an FDR < 0.05 and overall >were considered DEGs. Venn diagrams were generated to visualize the overlap between perturbed genes and DEGs.

### Pathway Analysis assessment: perturbed and differentially expressed genes

Gene set enrichment analysis of KEGG mouse pathways was conducted using the hypergeometric test. This analysis assessed both perturbed genes and DEGs. The pathways were ranked based on their p-values, and the top 10 pathways with the lowest p-values were selected.

### Validation using the Knockout Mouse Project Database

The KOMP database, accessible at the International Mouse Phenotyping Consortium website, serves as a valuable resource for researchers, offering comprehensive phenotypic data and genetic insights on a wide array of knockout mouse models. This platform facilitates the understanding of gene function and disease mechanisms and is instrumental in advancing the study of human diseases, including the identification of potential therapeutic targets.https://www.mousephenotype.org/understand/start-using-the-impc/impc-data-generation.

## Results

Given that scRNA-seq may fail to capture all disease-relevant genes, we focused our analysis on comparing and contrasting three gene categories: non-perturbed DEGs, perturbed non-DEGs, and perturbed DEGs, to evaluate their respective relevance to T2DM pathophysiology across its pathophysiological continuum to demonstrate that all three are essential to capture the full molecular spectrum of the disease. We selected the NZO/HlLtJ mouse strain as the mild and severe diabetic model as it displays signs of metabolic syndrome symptoms including morbid obesity, fasting hyperglycemia, hyperinsulinemia, insulin resistance, and hypercholesterolemia, and is a well-established mildly diabetic model^19–21^.

### Differentially expressed and perturbed **β** cells genes between prediabetic and healthy states

To identify prediabetic-state-specific perturbed genes, we performed ssNPA analysis using scRNA-seq of β cells isolated from healthy (C57BL/6J, standard diet) and prediabetic (C57BL/6J, HFHS diet) mice. In total, β cells exhibited 573 DEGs and 1,389 perturbed genes (**Figure 2A-B**). Of these, 289 genes were shared between DEGs and perturbed signatures (including 235 up-regulated and 54 down-regulated perturbed DEGs). We also identified 1,100 perturbed-only genes (perturbed but not DEGs) and 2,267 genes that were neither DEGs nor perturbed.

**Figure 2.**
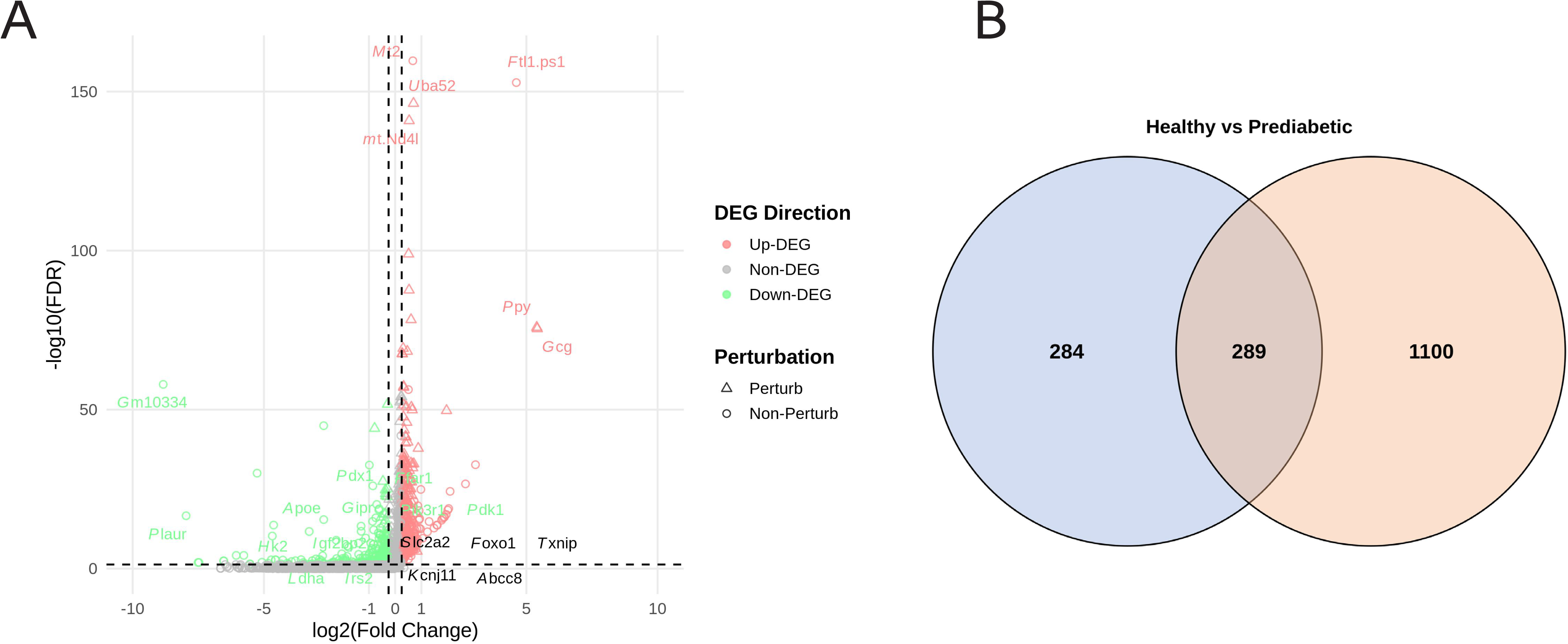
Differentially expressed and perturbed genes in β-cells from healthy and prediabetic mice. **A)** Volcano plot showing the relationship between gene expression changes (log□ fold change) and statistical significance (−log□□FDR) in β-cells from C57BL/6J mice fed a standard versus a high-fat, high-sucrose (HFHS) diet. Triangles denote perturbed genes, while circles represent non-perturbed genes; red and green indicate up- and downregulated DEGs, respectively. Several key metabolic regulators such as *Pdx1*, *Slc2a2*, *Foxo1*, and *Gcg* exhibit significant expression or perturbation differences between groups. **B)** Venn diagram depicting the overlap between DEGs (left) and perturbed-only genes (right). Of the total identified genes, 289 were shared between both categories, while 284 and 1,100 were unique to DEGs and perturbed-only sets, respectively, indicating that perturbation analysis reveals additional functionally altered genes beyond those identified by expression changes alone.

Among the non-perturbed DEGs, we identified several genes related to T2DM pathophysiology including *Igf2bp2, Irs2, Hk2, Ldha, Apoe, Klf2, Il1r2, Oas1, Oas2, Oas3, Rbp4, Slc2a3, Dusp4, Adm, and Inhba. Irs2 and Igf2bp2* are well-established T2DM susceptibility genes that regulate insulin and growth factor signaling *Irs2* acts as a major mediator of insulin and IGF signaling, ensuring proper β cell survival and glucose responsiveness^22,23^, whereas *Igf2bp2* stabilizes key metabolic transcripts that influence glucose homeostasis^9^. *Hk2* and *Ldha* are important glycolytic enzymes that help maintain cellular energy balance in β cells by regulating glucose phosphorylation and lactate production^24^. *Slc2a3* encodes a glucose transporter that enhances glucose uptake during periods of metabolic stress^25^, while *Rbp4* is an adipokine associated with insulin resistance and altered lipid metabolism in obesity and diabetes^26^.

Other genes in this group play significant roles in inflammation, stress response, and cellular remodeling. *Il1r2* acts as a decoy receptor that modulates interleukin-1–mediated inflammation in pancreatic β cells^27^*. Oas1*, *Oas2*, and *Oas3* are interferon-stimulated genes that may reflect immune activation contributing to β cell stress^28^. *Dusp4*^29^ *and Klf2*^30^ regulate transcriptional, and kinase pathways linked to oxidative and endoplasmic reticulum stress responses. *Apoe*^31^ *and Inhba*^32^ influence lipid handling and transforming growth factor signaling, both of which contribute to β cell dedifferentiation and senescence. *Adm* encodes adrenomedullin, a vasoprotective and cytoprotective peptide that supports islet perfusion and survival under metabolic load^33^. Together, these DEGs represent interconnected metabolic, inflammatory, and stress-adaptive mechanisms underlying the transition from β cell compensation to dysfunction in T2DM.

Among the perturbed non DEGs, we identified several genes with strong relevance to β cell function and T2DM pathophysiology, including *Glp1r, Kcnj11, Abcc8, Slc2a2, Foxo1, Txnip, Wfs1, Ptpn1* and *Dpp4*. *Glp1r* encodes the glucagon-like peptide 1 receptor, which mediates incretin signaling to enhance glucose-stimulated insulin secretion and promote β cell proliferation^10^. *Igf1r* regulates growth and survival signaling in β cells^34^. *Kcnj11* and *Abcc8* encode the subunits of the ATP sensitive potassium channel complex that is essential for glucose stimulated insulin secretion. Both have been associated with T2DM^35,36^. *Slc2a2* encodes GLUT2, the major glucose transporter that allows β cells to sense circulating glucose levels^37^. *Foxo1* integrates insulin and oxidative stress signaling and is key for maintaining β cell identity and proliferation^37^. *Txnip* connects oxidative stress to glucose toxicity by inhibiting thioredoxin^38^, while *Wfs1* protects the endoplasmic reticulum from stress and supports insulin folding and secretion^39^. *Ptpn1* encodes protein tyrosine phosphatase 1B, which downregulates insulin receptor signaling^40^, and *Dpp4* modulates incretin degradation, impacting insulin release^41^.

Additional perturbed non-DEGs such as *Pcsk1, Prlr, Rictor, Gcgr, Ptger* and *Nfe2l*., also play important roles in β cell metabolism and signaling. *Pcsk1* is involved in proinsulin processing^42^, while *Prlr* (prolactin receptor) supports β cell growth and functional adaptation during metabolic stress^43^. *Rictor*, a component of the mTORC2 complex, influences insulin signaling and cytoskeletal organization^44^. *Gcgr*^45^ and *Ptger3*^46^ mediate glucagon and prostaglandin signaling, both of which regulate calcium and cyclic AMP pathways that control insulin secretion. *Nfe2l2* encodes NRF2, a transcription factor that drives antioxidant defense and maintains redox homeostasis^47^. Together, these perturbed non-DEGs indicate early disruption of metabolic, oxidative, and signaling networks that compromise β cell adaptability before major transcriptional changes occur in diabetes progression.

Among the perturbed DEGs, we identified several genes closely linked to β cell signaling, metabolism, and stress adaptation, including *Pdx1, Ffar1, Gipr, Pik3r1, Pdk1, Slc2a1, Gcg, Ucn3, G6pc2, and Gdf15. Pdx1* is a master transcription factor for pancreatic β cell identity and insulin gene expression^11^. *Ffar1*^48^ and *Gipr* encode receptors that sense free fatty acids and incretins, respectively, enhancing glucose stimulated insulin secretion^49^. *Pik3r1*^50^ and *Pdk1*^51^ are core components of insulin signaling that regulate β cell survival and glucose metabolism. *Slc2a1* facilitates basal glucose uptake and compensatory glucose transport under stress^52^. *Gcg* encodes glucagon^53^, and *Ucn3* is a β cell secretory product that fine tunes paracrine signaling with α cells and is associated with T2DM development^54^. *G6pc2* participates in glucose cycling within β cells, influencing fasting glucose regulation^55^, while *Gdf15* acts as a stress responsive cytokine linked to metabolic adaptation and protection against glucolipotoxicity^56^.

### Differentially expressed and perturbed **β** cells genes between severely and prediabetic states

To identify diabetes-state-specific perturbed genes, we performed ssNPA analysis using scRNA-seq of β cells isolated from severely diabetic mice (NZO, HFHS diet) and prediabetic mice (C57BL/6J, standard diet). In total, β cells exhibited 1,289 DEGs (344 up-regulated and 945 down-regulated) and 413 perturbed genes (**Figure 3A-B**). Among these, 281 genes were shared between DEG and perturbed signatures (170 up-regulated perturbed DEGs and 111 down-regulated perturbed DEGs). We also identified 132 perturbed-only genes and 1,205 genes that were neither DEGs nor perturbed.

**Figure 3.**
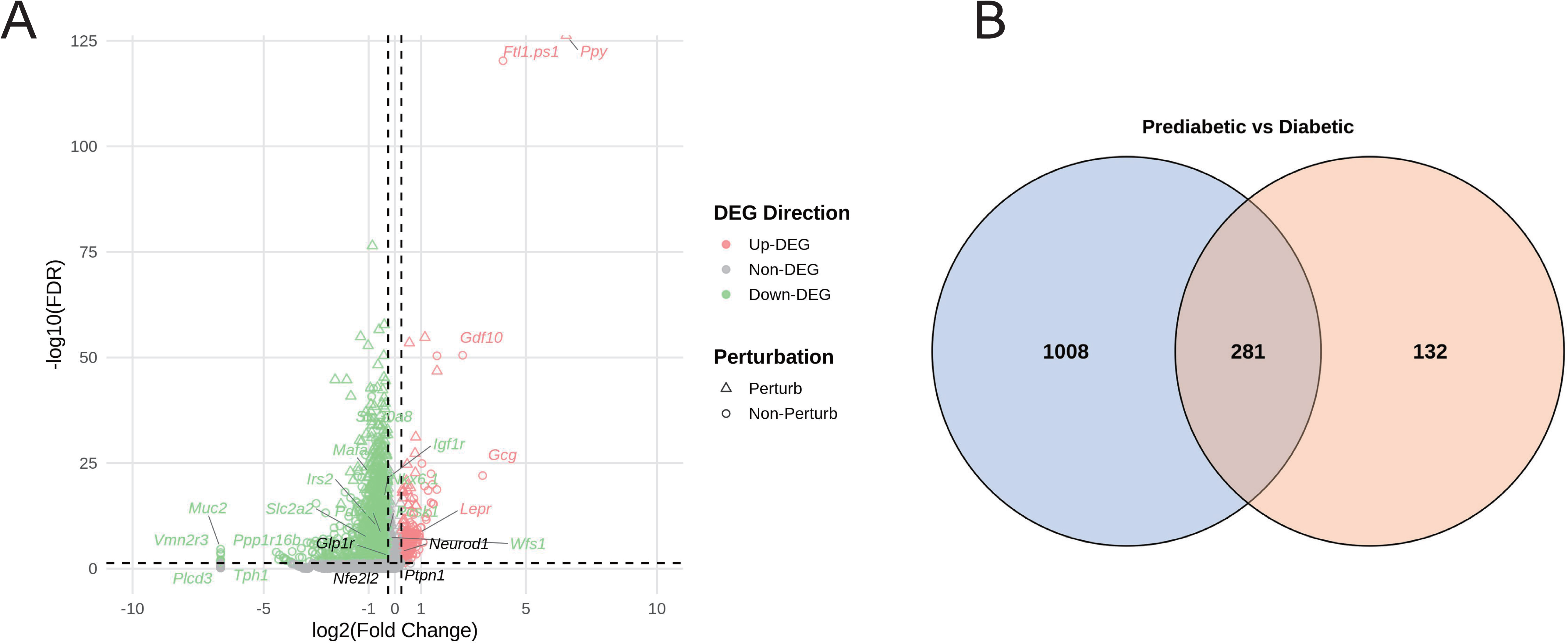
Differentially expressed and perturbed genes in β-cells from prediabetic and severely diabetic mice. **A)** Volcano plot illustrating the relationship between gene expression changes (log□fold change) and statistical significance (−log□□FDR) in β-cells from NZO/HILtJ mice fed a high-fat, high-sucrose (HFHS) diet compared to those on a standard diet. Triangles represent perturbed genes, while circles denote non-perturbed genes; red and green indicate up- and downregulated DEGs, respectively. Several critical β-cell and insulin signaling regulators, including *Glp1r*, *Slc2a2*, *Irs2*, *Wfs1*, and *Neurod1*, showed significant expression or perturbation differences between groups. **B)** Venn diagram showing the overlap between DEGs (left) and perturbed-only genes (right). Of the total identified genes, 281 were shared between both categories, while 1,008 and 132 were unique to DEGs and perturbed-only sets, respectively. This highlights that perturbation analysis uncovers additional network-level alterations beyond transcriptional changes detectable through differential expression alone.

Among these non-perturbed DEGs, we identified similar and additional several key regulators of β cell metabolism, insulin signaling, and nutrient sensing, including *Pdx1, Slc2a2, Irs2, Lepr, Fgf21, Ffar1, Pdk1, Pdk4, Pck2*, and *Hk2*. *Lepr*^57^ and *Fgf21*^58^ reflect endocrine crosstalk between adipose tissue, liver, and pancreatic islets, coordinating systemic energy balance and glucose homeostasis, while *Pck2* participates in gluconeogenic and cataplerotic flux, ensuring metabolic energy flexibility^59^. Additional genes such *as Gipr, Sst, Insig1, Cd36, Apoe, Rbp4, Ceacam1, Hhex, Maf,* and *Slc2a1* contribute to the integrated regulation of insulin secretion, lipid metabolism, and stress response. *Sst* encodes somatostatin, a paracrine inhibitor that modulates islet hormone release^60^. *Insig1* regulates lipid biosynthesis and endoplasmic reticulum homeostasis^61^. *Cd36*^62^, *Apoe*^31^, and *Rbp4*^26^ are involved in lipid and retinol transport, influencing insulin sensitivity and β cell lipid metabolism. *Ceacam1* promotes insulin clearance and maintains insulin receptor recycling^63^, while *Hhex*^64^ and *Maf*^65^ are transcription factors required for islet cell differentiation and function. Together, these genes form a tightly coordinated network that integrates glucose and lipid metabolism, insulin signaling, and intercellular communication to maintain β cell function and systemic glucose homeostasis.

Among the perturbed non-DEGs, several are critical for maintaining β cell identity, stress tolerance, and insulin signaling, including *Nkx6.1, Glp1r, Wfs1, Igf1r, Nfe2l2, Cebpb, Atf6, Relb, Bace2*, *Ppp1r15a* and *Cstb*. *Nkx6.1* is a core transcription factor required for β cell differentiation and insulin gene transcription, ensuring mature β cell function^66,67^. *Cebpb* and *Atf6* contribute to stress-adaptive transcriptional programs, particularly during ER stress and inflammatory signaling^68,69^. *Relb* functions in the noncanonical NF-κB pathway, mediating inflammatory tolerance^70^, while *Bace2* modulates proinsulin processing and insulin granule maturation^71^. *Ppp1r15a*, also known as GPR43 is a key regulator of the integrated stress response that promotes recovery from ER stress through dephosphorylation of eIF2α^72^. *Cstb,* a potential exerkine^73^, regulates lysosomal protease activity, protecting cells from proteotoxic stress and may contribute to T2DM^74^. Collectively, these genes represent a cohesive stress-adaptive and survival network that safeguards β cell function in the face of metabolic overload and inflammatory stress characteristic of diabetes progression.

Among the perturbed DEGs, several are central regulators of β cell differentiation, glucose sensing, and insulin secretion, including *Mafa, Neurod1, Pcsk1, Slc30a8, Ptpn1, Ucp2, Ddit3, Socs3, Abca1*, and *C2cd4b*. *Mafa* and *Neurod1* are key transcription factors essential for maintaining mature β cell identity and activating insulin gene expression. In particular, *MAFA*-associated missense mutations have been implicated in familial insulinomatosis and diabetes. *Slc30a8* encodes the zinc transporter ZnT8, crucial for insulin crystallization and storage within secretory granules and its mutations are linked with T2DM^75^. *Ptpn1* negatively regulates insulin receptor signaling, and its overactivity contributes to insulin resistance and β cell stress^76^. *Ucp2* modulates mitochondrial coupling and reactive oxygen species generation, fine-tuning insulin secretion efficiency^77^. *Ddit3* (CHOP) is a marker of ER stress and a key mediator of β cell apoptosis during chronic metabolic overload^78^. *Socs3* suppresses cytokine and insulin signaling under inflammatory conditions^79^, while *Abca1* regulates cholesterol efflux, preserving membrane integrity and insulin granule function^80^. *C2cd4b* has been genetically linked to T2DM susceptibility, possibly influencing calcium-mediated signaling in β cells^81^.

### Differentially expressed and perturbed **β** cells genes between severely and mildly diabetic sages

To identify diabetes-state–specific perturbed genes, we performed ssNPA analysis using scRNA-seq of β cells isolated from severely diabetic mice (NZO, HFHS diet) and midly diabetic mice (NZO, standard diet). In total, β cells exhibited 1,293 DEGs (1,371 down-regulated and 162 up-regulated) and 966 perturbed genes (**Figure 4A-B**). Among these, 637 genes were shared between DEG and perturbed signatures (566 down-regulated perturbed DEGs and 71 up-regulated perturbed DEGs). We also identified 329 perturbed-only genes (perturbed but not differentially expressed) and 1,946 genes that were neither perturbed nor differentially expressed.

**Figure 4.**
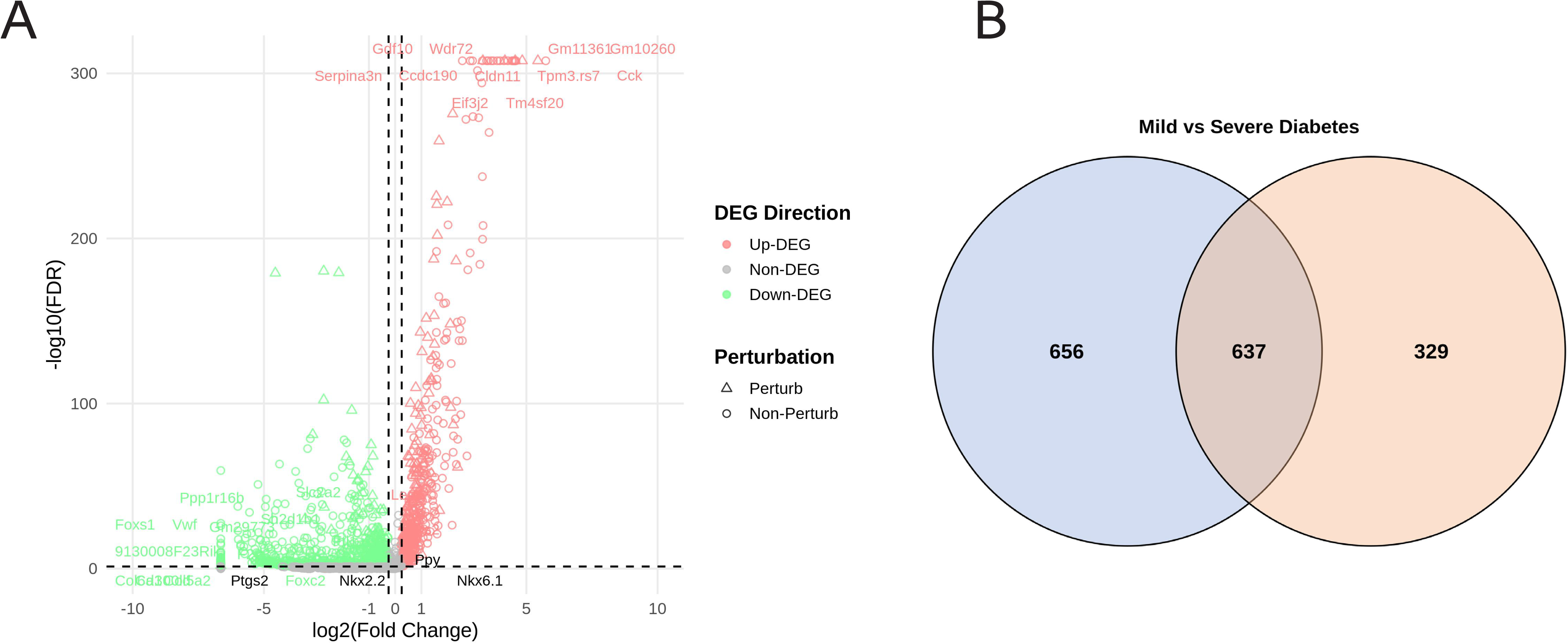
Differentially expressed and perturbed genes in β-cells from mild and severe diabetic mice. **A)** Volcano plot showing the relationship between gene expression changes (log□ fold change) and statistical significance (−log□□FDR) in β-cells from NZO/HILtJ mice with mild versus severe diabetes. Triangles denote perturbed genes, while circles represent non-perturbed genes; red and green indicate up- and downregulated DEGs, respectively. Several metabolic and β-cell transcriptional regulators, including *Pparg*, *Foxa2*, *Slc2a2*, *Lepr*, and *Gcg*, exhibited marked differences in expression or perturbation between disease stages. **B)** Venn diagram showing the overlap between DEGs (left) and perturbed-only genes (right). A total of 637 genes were shared between both categories, while 656 and 329 were unique to DEGs and perturbed-only sets, respectively. These findings suggest that as diabetes severity increases, both transcriptional dysregulation and network-level perturbations become more extensive, reflecting progressive β-cell dysfunction.

Among these genes, several are central to insulin signaling, glucose metabolism, and energy regulation, including *Pparg*, *Ppargc1a*, *Irs2*, *Slc2a2*, *Lepr*, *Dpp4*, *Rbp4*, *Camk1d*, *Pdk1*, and *Gcg*. *Pparg* and *Ppargc1a* coordinate lipid and glucose metabolism, promoting insulin sensitivity and mitochondrial biogenesis in peripheral tissues^82^. *Camk1d* regulates calcium-dependent signaling pathways tied to insulin secretion and is associated with increased risk for T2DM^83^. Additional genes such as *Hk1*, *Ldha*, *Ffar4*, *Ffar2*, *C2cd4a*, *Sec16b*, *Ksr2*, *Cdk4*, *Adm*, and *Abcc9* contribute to the metabolic flexibility and energy homeostasis of β cells. *Hk1* initiates glycolysis by phosphorylating glucose and its disruption is implicated in T2DM^84^. *C2cd4a* has been genetically linked to T2DM and likely regulates calcium-mediated exocytosis in β cells^85^. *Sec16b* is involved in endoplasmic reticulum export and secretory pathway maintenance and has been recently identified as a key regulator of glucose homeostasis^86^. *Ksr2* acts as a scaffold protein regulating AMPK and ERK signaling, linking cellular energy status with metabolic output and is involved in T2DM pathogenesis^87^. *Cdk4* promotes β cell proliferation and adaptive expansion during insulin resistance^88^. *Abcc9* encodes the SUR2 subunit of ATP-sensitive potassium channels, influencing membrane excitability and insulin release, and along with *Abcc8,* contribute to T2DM^89^. Collectively, these genes form a network that integrates glucose and lipid metabolism, hormonal signaling, and adaptive stress responses that are fundamental to β cell compensation in diabetes.

Among the perturbed non-DEGs, several are key regulators of pancreatic lineage specification, mitochondrial metabolism, and insulin signaling, including *Mt.nd1*, *Glp1r*, *Nkx2.2*, *Foxa2*, *Ndufaf2*, *Atp1b1* and *Fosl2* . *Mt.nd1* encodes a mitochondrial NADH dehydrogenase subunit essential for oxidative phosphorylation and ATP generation, supporting the high energy demands of insulin secretion^90^. *Nkx2.2* and *Foxa2* are core transcription factors governing pancreatic development, β cell differentiation, and maintenance of glucose-sensing machinery^91,92^. *Ndufaf2* participates in mitochondrial complex I assembly, ensuring efficient electron transport and cellular respiration^93^. *Atp1b1* encodes a subunit of Na /K -ATPase, which maintains membrane potential and ion gradients critical for insulin vesicle exocytosis^94^. *Fosl2* is part of the AP-1 transcriptional complex that modulates cellular growth and metabolic gene expression^95^.

Among the perturbed DEGs, several play critical roles in mitochondrial metabolism, β cell differentiation, and glucose utilization, including *Mt.nd2*, *Nkx6.1*, *H2.t22*, *Mt.atp8* and *Pappa2*. *Mt.nd2* and *Mt.atp8* encode mitochondrial components of the electron transport chain, essential for ATP production required for insulin exocytosis^96,97^. *H2.t2 is* an MHC-related genes, potentially linking immune modulation with metabolic stress responses in β cells^98^. *Pappa2* modulates local IGF signaling by cleaving IGF-binding proteins, enhancing insulin sensitivity and growth factor bioavailability^99^.

### Comparative overlap of differentially expressed and perturbed genes across the diabetes continuum

To further illustrate that DEGs and perturbed genes capture both unique and shared molecular signatures across the T2DM continuum, we compared the number of overlapping genes across all three conditions and between each condition (**Figure 5A–F**, see **Supplementary Tables 1–4** for the full list of genes).

**Figure 5.**
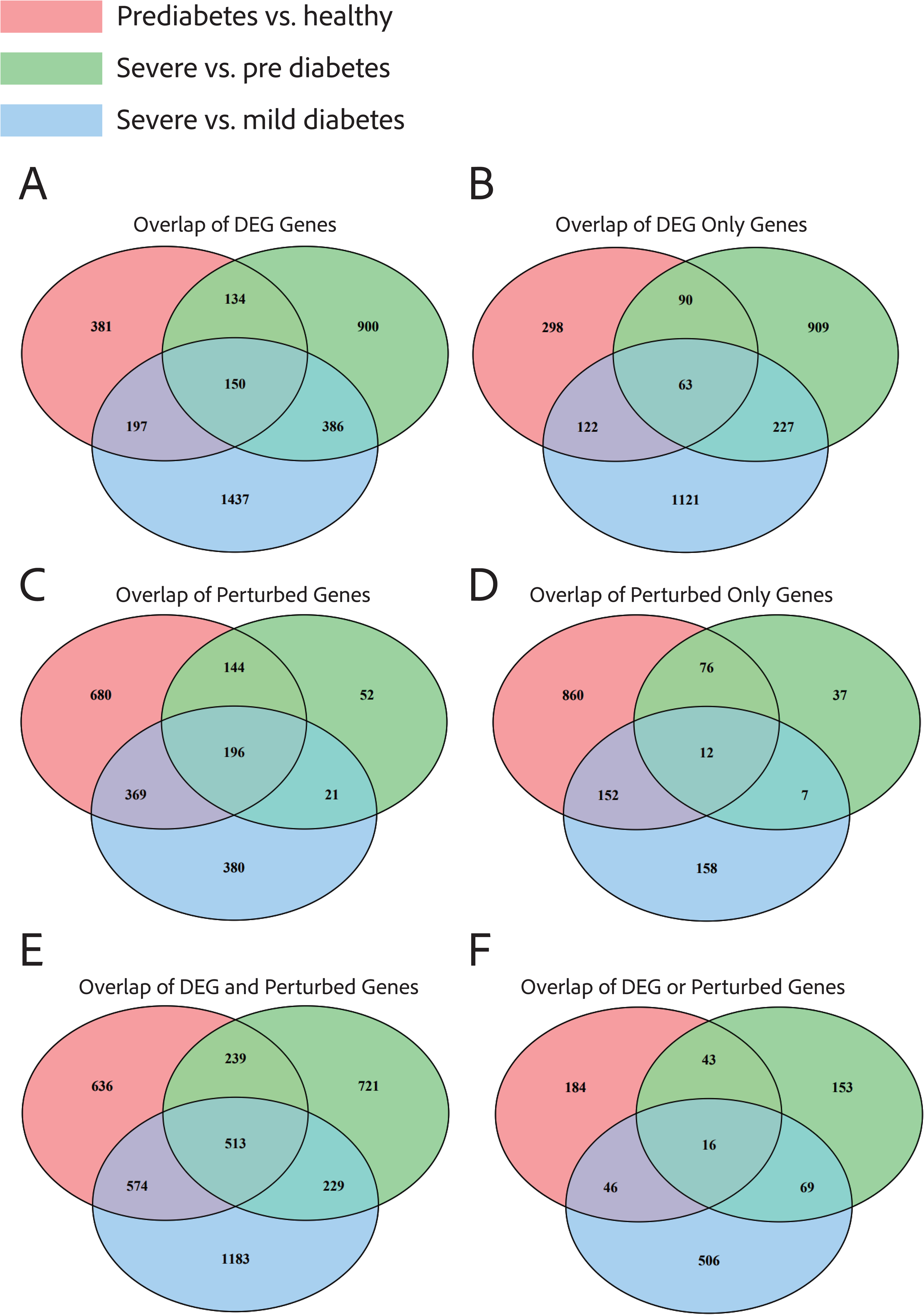
Overlap of DEGs and perturbed genes across stages of the T2DM continuum. Venn diagrams show the overlap in differentially expressed and perturbed genes identified across three comparisons: prediabetes versus healthy (red), severe versus prediabetes (green), and severe versus mild diabetes (blue). **A)** Overlap of all DEGs across conditions, revealing both shared and stage-specific transcriptional changes. **B)** Overlap of DEGs only, indicating a limited set of consistently altered genes across all comparisons. **C)** Overlap of all perturbed genes, showing broader network-level disruptions shared between disease stages. **D)** Overlap of perturbed-only genes, highlighting 12 genes shared across all three conditions that were not captured by differential expression alone. **E)** Overlap between DEGs and perturbed genes, demonstrating partial convergence between expression-level and network-level changes. **F)** Overlap of DEGs or perturbed genes, showing the combined molecular landscape across the T2DM continuum. Collectively, these results illustrate that DEG and perturbation analyses capture both distinct and overlapping molecular features underlying progressive T2DM pathophysiology.

We identified 63 non-perturbed DEGs shared across all three comparisons (**Figure 5B**), including *Irs2*. *Irs2* plays a central role in insulin signaling, transmitting signals from the insulin and IGF-1 receptors to downstream pathways like PI3K-Akt that regulate glucose uptake, β-cell survival, and hepatic glucose production. Impaired *Irs2* expression or phosphorylation is a well-established contributor to insulin resistance and β-cell failure, two key hallmarks of T2DM. Mouse models lacking *Irs2* develop severe diabetes due to defective β-cell compensation, confirming its causal role in maintaining glucose homeostasis^100^.

We identified 12 perturbed-only genes shared across all three comparisons (**Figure 5D**), including *Glp1r,* a clinically relevant gene encoding the glucagon-like peptide-1 receptor. Glp1r is a key therapeutic target in T2DM, as it mediates insulin secretion and is the basis for GLP-1 receptor agonist drugs used to improve glycemic control and promote weight loss^10^. Interestingly, despite its strong association with T2DM, Glp1r was not detected as a DEG in the scRNA-seq data, suggesting that network-level perturbation analysis can uncover functionally important regulatory changes that are not apparent from expression-level differences alone.

We identified 16 perturbed DEGs shared across all three comparisons (**Figure 5F)**. These included *Serping1*, which encodes C1 esterase inhibitor, a key regulator of the complement and coagulation cascades, which have been implicated in metabolic inflammation, endothelial dysfunction, and insulin resistance in T2DM. Consistent with prior microarray analyses^101^, *Serping1* was identified among the core genes enriched in pathways related to pancreatic secretion and immune activation, suggesting that its dysregulation may contribute to β-cell stress and impaired insulin signaling.

Together, these findings demonstrate that integrating DEG and perturbation analyses provides a more comprehensive view of T2DM pathophysiology. While DEGs highlight transcriptional shifts directly measurable at the expression level, perturbation-based analysis captures regulatory and signaling alterations that may occur without major expression changes. The overlap and divergence between these layers reveal distinct yet complementary molecular mechanisms across the T2DM continuum.

### Validation of perturbed genes using the KOMP database

To corroborate the perturbed genes identified with the meta-ssNPA analysis, we employed the KOMP database and generated visual representations for the genes previously linked to T2DM. The *in vivo* mouse reference data for validation of genes was generated by the Knock-out Mouse Project-KOMP (www.mousephenotype.org)^102,103^. The perturbed genes selected for validation were based on their response to a glucose tolerance test, the standard method to diagnose diabetic or obese phenotypes in both preclinical and clinical research. We selectively chose three genes that demonstrated improved glucose tolerance/intolerance based on their area under curve profiles (AUC) and have been associated with T2DM: *Glp1r, Abcc8, and Kcnj11*.

Across all three knockout models, glucose tolerance tests revealed significant impairments in glucose clearance compared to wild-type controls, consistent with diabetic phenotypes (**Figure 6A-C**). These findings validate the computationally predicted perturbations, as all three genes are established regulators of pancreatic β-cell excitability and insulin secretion. Loss of *Glp1r* impairs incretin-mediated insulin release^10^, while *Abcc8* and *Kcnj11* encode subunits of the ATP-sensitive potassium (K_ATP) channel essential for insulin granule exocytosis^104–106^.

**Figure 6.**
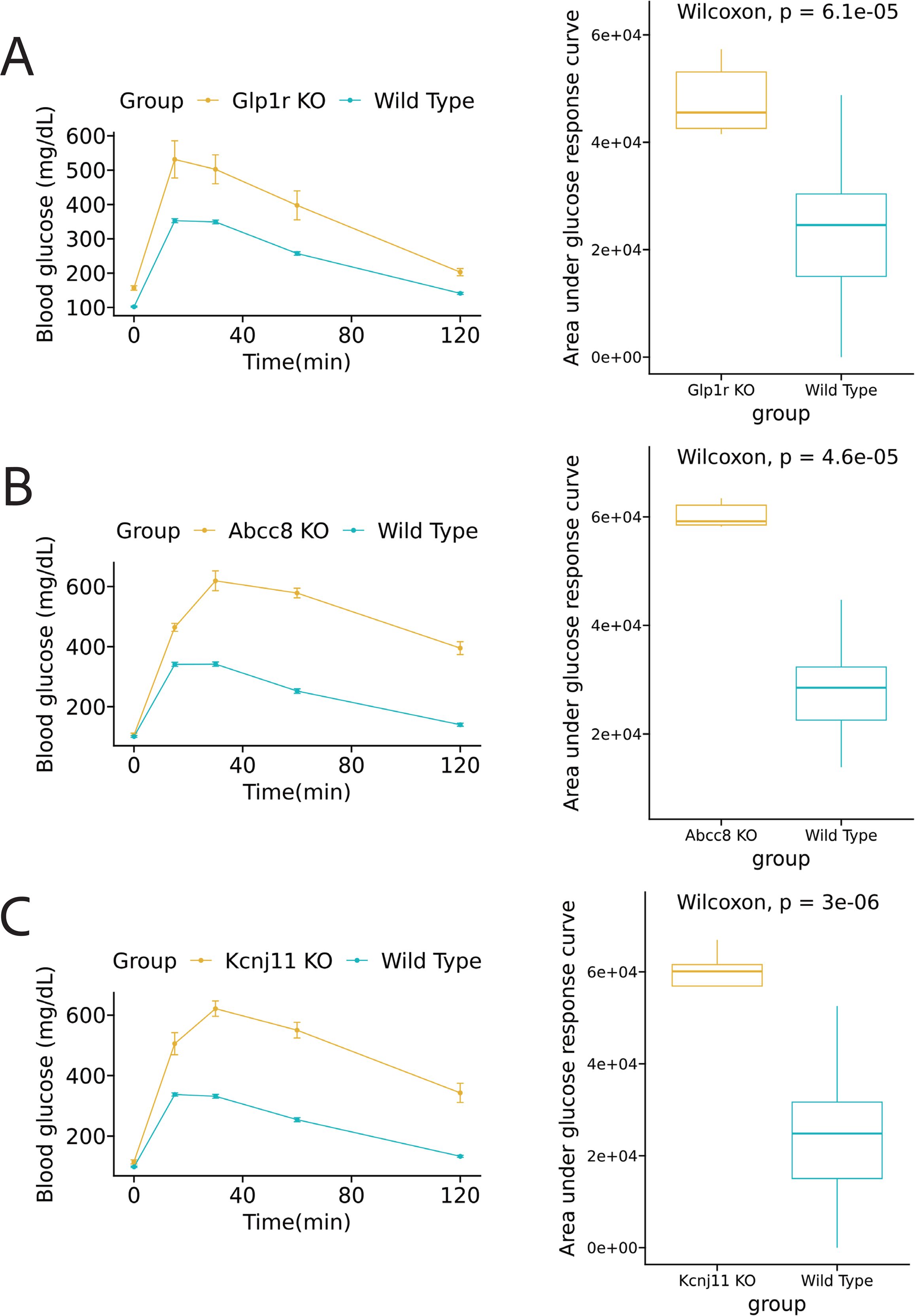
In vivo validation of perturbed genes using the Knockout Mouse Phenotyping (KOMP) database. Glucose tolerance test (GTT) data from knockout (KO) mice were analyzed to validate genes identified as perturbed in β-cells. **(A–C)** Time-course plots show blood glucose levels following glucose administration in *Glp1r*, *Abcc8*, and *Kcnj11* KO mice (orange) compared with wild-type controls (blue). KO mice exhibited significantly impaired glucose clearance, reflected by elevated glucose levels over time. **(A–C, right)** Box plots of the area under the glucose response curve (AUC) further confirm reduced glucose tolerance in KO mice relative to controls, with highly significant Wilcoxon p-values. These findings corroborate the role of *Glp1r*, *Abcc8*, and *Kcnj11* as key regulators of β-cell function and glucose homeostasis.

## Discussion

Several studies have utilized gene expression data to construct causal networks^107–109^. Additionally, researchers have identified gene features that exhibit strong predictive power for specific phenotypes^110–112^. In this context, the innovative approach of ssNPA assesses how the gene network of a set of control samples is perturbed when presented with a new query sample. The underlying rationale of ssNPA is based on the notion that, in many diseases, an observed phenotype may arise from alterations in different components of the ’healthy’ gene network. The ssNPA framework offers several distinct advantages over traditional approaches. Firstly, it enables the inference of topological relationships among genes within the causal network providing valuable insights into the interconnections and regulatory dynamics among genes. Secondly, the data-driven nature of the network allows for the discovery of novel information enhancing our understanding of the complex mechanisms underlying the disease. Thirdly, ssNPA does not rely on prior pathway knowledge, making it a valuable tool for investigating gene perturbations in a hypothesis-free manner. In comparison, prior studies on T2DM primarily relied on DEG analysis and genome-wide association studies (GWAS), which lack the aforementioned advantages of the ssNPA framework.

T2DM is a chronic metabolic disorder characterized by heterogeneity and polygenic traits. The genetic bases of T2DM are poorly understood, highlighting the need of approaches that can help investigate the genes and associated signaling pathways contributing to its onset. To this end, we constructed a transcriptional network that explored perturbed genes, hub genes, and associated pathways and performed validation using the KOMP database. The ssNPA enabled the identification of several genes already established in the diabetes context while also unveiling novel genes previously unrecognized in relation to diabetes. In our comparison between C57BL/6J mice on a regular chow diet and those on a HFHS diet, the analysis highlighted a significant number of perturbed genes. This observation is particularly noteworthy as there is a limited report of genes during the *prediabetic* state, emphasizing the importance of these findings in understanding new-onset diabetes/prediabetes. The key findings in this comparison included genes, such as *Glp1r* and *Abcc8*, which we subsequently validated using the KOMP database.

SsNPA is a sophisticated method designed to assess perturbations in gene networks at the level of individual samples. Instead of focusing on isolated genes, ssNPA delves into the broader landscape of interconnected gene networks. The methodology begins by inferring a global gene network, utilizing causal graph learning derived from a set of reference samples. Upon the introduction of a new sample, ssNPA calculates the degree of deviation of this sample from the established reference network at every gene point. This approach furnishes in-depth information regarding the topology of network perturbations. By generating a perturbation feature vector, i.e. perturbance score, ssNPA allows for the classification or clustering of samples, which can be instrumental in distinguishing between cell types, disease subtypes, or any other biological distinctions under investigation. This tool provides an advanced perspective on how various perturbations, such as environmental changes, drug treatments, or genetic mutations, affect the broader dynamics of gene networks. SsNPA and differential gene expression analysis, while both employed in the domain of genomics, serve distinctly different analytical purposes. Differential gene expression analysis aims to identify individual genes that exhibit statistically significant differences in expression between two or more conditions, such as healthy versus diseased states. Its output is often a list of upregulated or downregulated genes. In contrast, ssNPA focuses on assessing gene network perturbations in individual samples. Instead of concentrating on the behavior of single genes, ssNPA evaluates how perturbations, such as mutations or drug treatments, influence entire gene networks or pathways. It provides insights into deviations from a reference network, shedding light on both the magnitude and topology of network perturbations. While differential gene expression gives a granular view of specific genes’ behavior, ssNPA offers a holistic perspective on how interconnected gene networks respond to various conditions.

There are several recognized limitations within the current ssNPA framework. Primarily, the framework exhibits suboptimal performance when applied to single-cell data. Such data often present sparsity challenges, characterized by numerous genes that register zero expressions across a multitude of cells. This phenomenon disrupts the linear model’s foundational assumption upon which ssNPA operates, resulting in diminished prediction accuracy. A proposed solution to this challenge, introduced by Baran and Bercovich^16^, involves the innovative ’meta-cell’ concept. By randomly selecting multiple cells of identical cell types and computing the mean gene expressions, a novel meta-cell sample is generated. This methodology retains the relative gene expression hierarchy within the cells, ensuring genes with higher expressions remain dominant, and those with lower expressions continue to be subdued in the meta-cells, while concurrently mitigating the zero-expression issue. A secondary concern pertains to the criteria ssNPA employs to discern Perturbed genes. In its original design, a gene is classified as “Perturbed” provided its average Perturbance Score exceeds that of the control group. This method, however, overlooks the potential influence of random noise on the Perturbance Score, which can inadvertently result in False Positive outcomes. To enhance precision, we integrated a statistical approach, employing the Wilcoxon test as elucidated by Li et al.^17^. By contrasting the Perturbance Scores of both test and control groups and setting a FDR threshold of less than 0.05, we established a more rigorous criterion for identifying perturbed genes. For this analysis, a one-sided Wilcoxon test was executed, operating under the alternative hypothesis that the control group’s Perturbance Score is inferior to that of the test group. The KOMP validation process is designed to be broad and encompasses multiple cell types, rather than being specific to any single cell type. This approach allows for a more generalizable understanding of gene function across different biological contexts.

## Conclusion

In conclusion, our study employed meta-ssNPA, an innovative combination of meta-cell transcriptome analysis with the ssNPA framework, to investigate gene perturbations in various conditions. We identified genes that are perturbed in the context of T2DM and metabolic disorders, shedding light on potential therapeutic targets. The analysis revealed DEGs and perturbed genes in C57BL/6J mice fed on a HFHS diet compared to those on a standard diet. Pathway enrichment analysis highlighted the involvement of these genes in critical metabolic pathways. Moreover, we identified known T2DM genes such as *Glp1r*, *Igf1r*, and *Zbtb20*. which have previously been linked to metabolic traits and insulin regulation. Novel genes such as *Bpifc*, *Itga11*, *P2rx2*, *Tnip1* and *Khl32* are also discovered by the analysis and validated through KOMP database. To validate our findings, we leveraged the KOMP databasex knockout and characterize all protein-coding genes in the mouse genome, which corroborated the identified perturbed genes obtained from the meta-ssNPA analysis. Additionally, we discovered novel genes that exhibited perturbations despite not being DEGs and not directly associated with T2DM.

## Acknowledgments

The authors thank the Knock-out Mouse Project (KOMP) and members of Li Lab for discussions. The authors thank The Jackson Laboratory Computational Sciences and Research IT teams for technical support. Research reported in this publication was partially supported by The Jackson Laboratory Cube Initiatives. S. Li was supported by startup funds from The Jackson Laboratory and by the National Cancer Institute of the National Institutes of Health under Award Number P30CA034196. S. Li is also supported by the National Institute of General Medical Sciences of the National Institutes of Health under Award Number R35GM133562, by the National Human Genomic Research Institute of the National Institutes of Health under Award Number U01HG013175, by the National Cancer Institute of the National Institutes of Health under Award Number U01CA271830 and U01CA271830-03S1, by the NIH Common Fund under Award Number U54AG079753, and by the National Institute of Aging of the National Institutes of Health under Award Number R56AG071766.

